# Whole-genome sequencing and single acute toxicity of heavy metal to *Photobacterium kishitanii* FJ21

**DOI:** 10.1101/2022.09.20.508755

**Authors:** Shuzheng Yin, Zilong Yi, Jia Liu, Gang Liu, Jun Fang

## Abstract

In this study, the growth morphology of FJ21 strain was observed, and its 16S rRNA and whole genome were sequenced. Then, related software was used to make genome assembly, gene structure and function annotation, genome phylogenetic tree analysis, genome collinearity analysis and prediction of secondary metabolic gene cluster analysis. Finally, the single acute toxicity of five heavy metals to FJ21 strain was detected. There were *lux*C, *lux*D, *lux*A, *lux*B, *lux*F, *lux*E and *lux*G genes in FJ21, and the protein encoded by *lux* operon had certain hydrophilicity. The genome of this strain FJ21 contains a chromosome with a total length of 4853277bp and a GC content of 39.23%. The genome of FJ21 was compared with that of *Photobacterium kishitanii* ATCCBAA-1194, *Photobacterium phosphoreum* JCM21184, *Photobacterium aquimaris* LC2-065, *Photobacterium malacitanum* CECT9190, and *Photobacterium carnosum* TMW 2.2021. The average nucleotide identity(ANI), tetra nucleotide signatures (Tetra), comparative genome, and phylogenetic analysis proposed that FJ21 is a strain of *Photobacterium kishitanii*. In the acute toxicity test, the toxicity of heavy metals to the strain FJ21 is Pb(NO_3_)_2_ > ZnSO_4_·7H_2_O > CdCl_2_·2.5H_2_O > CuSO_4_·5H_2_O > K_2_Cr_2_O_7_.

## 1. Introduction

Luminescent bacteria are a group of Gram-negative bacteria that can emit blue-green fluorescence under normal conditions, and live in the marine environment mainly in the form of free organisms or parasites. The bioluminescence of luminescent bacteria is regulated by the enzyme-catalyzed reaction encoded by the lux operon. The *lux*A and *lux*B genes encode the α and β subunits of luciferase, respectively. *lux*C, *lux*D a,nd *lux*E constitute the fatty acid reductase complex, responsible for the synthesis of long Aldehyde substrate, luxG encodes flavin reductase[1]. Conserved genes *lux*C, *lux*D, *lux*A, *lux*B, *lux*E, and *lux*G exist in all luminescent bacteria that have been discovered[2], in addition, there are other genes such as *lux*I, *lux*R, and *lux*F[3]. Although luminescent bacteria have the same *lux* operon, these bacteria show great differences in characteristics such as growth behavior, luminescence intensity, or bioluminescence regulation[4]. The luminescence process of luminescent bacteriis oxidizedze FMNH_2_ and RCHO to FMN and RCOOH under the catalysis of intracellular specific luciferase and the participation of molecular oxyen, and at the same time release blue-green light. The luminescence reaction is as follows:

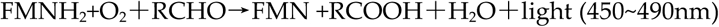

Luminescent bacteria can not only act as biosensors[5] but also produce antibacterial compounds[6], lipase[7], asparagine[8], and esterase[9]. Due to the luminescent bacteria method has the advantages of high sensitivity, simple processing, rapid response, and real-time monitoring, it has been widely used in the monitoring of water toxicity and environmental pollutants[10], and the acute and chronic toxicity tests of heavy metal mixtures[11]. At present, the luminescent bacteria commonly used in water quality and environmental monitoring are *Photobacterium phosphoreum, Vibrio fischeri*, and *Vibrio Qinghaiensis*[12]. The bright luminescence is usually used in the national standard GB/T15441-1995 for the determination of acute toxicity of water quality. The basic principle of luminescent bacteria for acute toxicity detection is that the luminescence process is easily affected. As long as the respiration or physiological process of bacteria is disturbed, the luminescence intensity of the bacteria will change[13]. In recent years, with the development of high-throughput sequencing technology, many microorganisms have completed genome sequencing. Whole-genome sequencing is an important foundation of microbial molecular mechanism research and development.

In the study, 16S rRNA, genome phylogenetic tree analyses, comparative genomics, average nucleotide identity (ANI), and tetra nucleotide signatures (Tetra) were used to clarify the strain FJ21 to *Photobacterium kishitanii*. In addition, the single acute toxicity of heavy metals to strain FJ21 was detected, which provided some data support for ecological risk assessment of heavy metal pollution.

## 2. Materials and Methods

### 2.1. The growth morphology, 16S rRNA and lux operator analysis

Beef extract, tryptone, sodium chloride, potassium dihydrogen phosphate, disodium hydrogen phosphate, glycerol, and agar. After the glycerol-preserved strain was cultured at 25°C for 24h, its morphological characteristics (size, shape, transparency, color, etc.) were observed.

Bacterial 16S rRNA universal primer was selected, and the PCR products were sent to Sangong Biotech (Shanghai) Co., Ltd. for sequencing, and the sequencing results were blast analyzed in the GenBank database. At the same time, the phylogenetic tree was constructed with mega-x software.

Prediction of protein secondary structure by protein analysis online software SWISS-MODEL.

### 2.2. Genome Sequencing and Assembly

Nanopore sequencing technology[14] was used to complete the genome scanning and sequencing of the strain. Firstly, high-quality DNA was extracted with a Qiagen kit, anan d ID library was constructed. The DNA was sequencea d by single moleculethe using Oxford Nanopore Technology sequencing instrument PromethION to obtain the original sequencing data. After quality control of the obtained sequencing data, the whole genome scanning of the strain was completed by bioinformatics analysis.

Assembly: the three-generation data after quality control were assembled with flye(parameter : --plasmids --nano-raw), corrected with racon(parameter : default) combined with three-generation sequencing data, and corrected with pilon[15] or NextPolish[16] (parameter : default) combined with two-generation sequencing data. The corrected genome uses its script to detect whether the loop is formed. After removing redundant loops, the origin of the sequence is moved to the replication initiation site of the genome by the circulator[17] (parameter : fixstart), to obtain the final genome sequence.

### 2.3. Genome Structure and Function Annotation

Genome structure prediction includes coding gene prediction, non-coding gene prediction, CRISPR prediction, and gene island prediction. The coding gene was predicted by prodigal[18] (parameter : -p None -g 11), and the complete CDS was retained. In the prediction of non-coding genes, RNAmmer[19] (parameter:-kingdom bac) and tRNAscan-SE2.0[20] (parameter:-B -I -m lsu,ssu,tsu) software were used to predict rRNA and tRNA in the genome, respectively. Other non-coding RNAs (ncRNAs) were predicted by the Infernal 1.1 [21] (parameter : --cut_ga --rfam --nohmmonly) search Pfam 13.0 database [22], and the predicted length was greater than 80 % of the sequence length in the database. CRISPR(parameter : default) was predicted by minced, and gene island was predicted by Islander[23] (parameter r:default).

After extracting the genome-coded proteins, InterProScan 5[24] was used for annotation, and the annotation information of TIGRFAMs[25], Pfam[26], and GO[27]databases were extracted. Blastp was used to compare the encoded proteins to KEGG[28], Refseq[29], and COG[30] databases, and the best results with a coverage of more than 30 % were retained as annotation results. The interaction genes between pathogen and host were annotated bythe PHI database, and the antibiotic resistance genes were annotatethe d by CARD database.

### 2.4. Phylogenetic tree Analysis and Sequence-based method for species identification

MEGA7.0 software was used to analyze the strain FJ21 and construct a phylogenetic tree by the Neighbor-Joining method.

The According to the recommended cut-off values for species determination (<95% for ANIb/ANIm and <0.989 for Tetra) [31,32], the calculation of average nucleotide identity based on BLAST(ANIb)/MUMmer(ANIm) and the correlation indexes of tetra nucleotide signatures (Tetra) were conducted using JspeciesWS (http://jspecies.ribohost.com/jspeciesws/#Analyse) [33,34].

### 2.5. Genome collinearity analysis

The whole-genome sequencing of strain FJ21 was analyzed by collinearity with other genome sequences with a similarity of 95 % in the NCBI database. MUMmer (version 3.23) was used to quickly compare the genomes of strain FJ21 and five closely related strains (*Photobacterium kishitanii* ATCCBAA-1194, *Photobacterium phosphoreum* JCM21184, *Photobacterium aquimaris* LC2-065, *Photobacterium malacitanum* CECT9190, and *Photobacterium carnosum* TMW 2.2021). Visualize each Contig of the genome using a brown box in the ggplot2 package in R language. Yellow lines between genomes represent Colinear and blue lines between genomes represent Inversion.

### 2.6. Prediction of secondary metabolite gene cluster

The assembled genome was analyzed by using antiSMASH version 6.0.0, and the parameters were selected from taxon bacteria.

### 2.7. For detection of heavy metal toxicity

The slant test-tube strains were eluted with 2ml of 3% NaCl solution, and after fully shaking, 200μL was taken into 50 ml of liquid culture medium, and cultured at 25 °C for 20-22h. Take 200μL bacterial solution into 40 ml of 3% NaCl solution and stir for 40 min for detection.

Dilute the solution of heavy metals(ZnSO4·7H_2_O, CuSO_4_·5H_2_O, Pb(NO_3_)_2_, CdCl_2_·2.5H_2_O, K_2_Cr_2_O_7_) to be measured with 3%NaCl into a series of concentration gradients. Add 0.1mL of the balanced bacterial solution into each sample tube, then take 1.9mL of the substance to be tested, and take 1.9mL of 3%NaCl as a blank. After 15min exposure, the luminous intensity was measured by portable Lux-T020 toxicity analyzer, and the dose-effect curve was drawn to obtain the half effect concentration (EC_50_).

## 3. Results and Discussion

### 3.1. The growth morphology,16S rRNA and lux operator analysis

#### 3.1.1. Growth morphology of strain

After the strain is cultured for 24 hours, the colony shape is round, slightly raised, smooth, milky white and short rod, which emits bright blue-green light in the dark, as shown in Fig 1.

**Figure 1.**
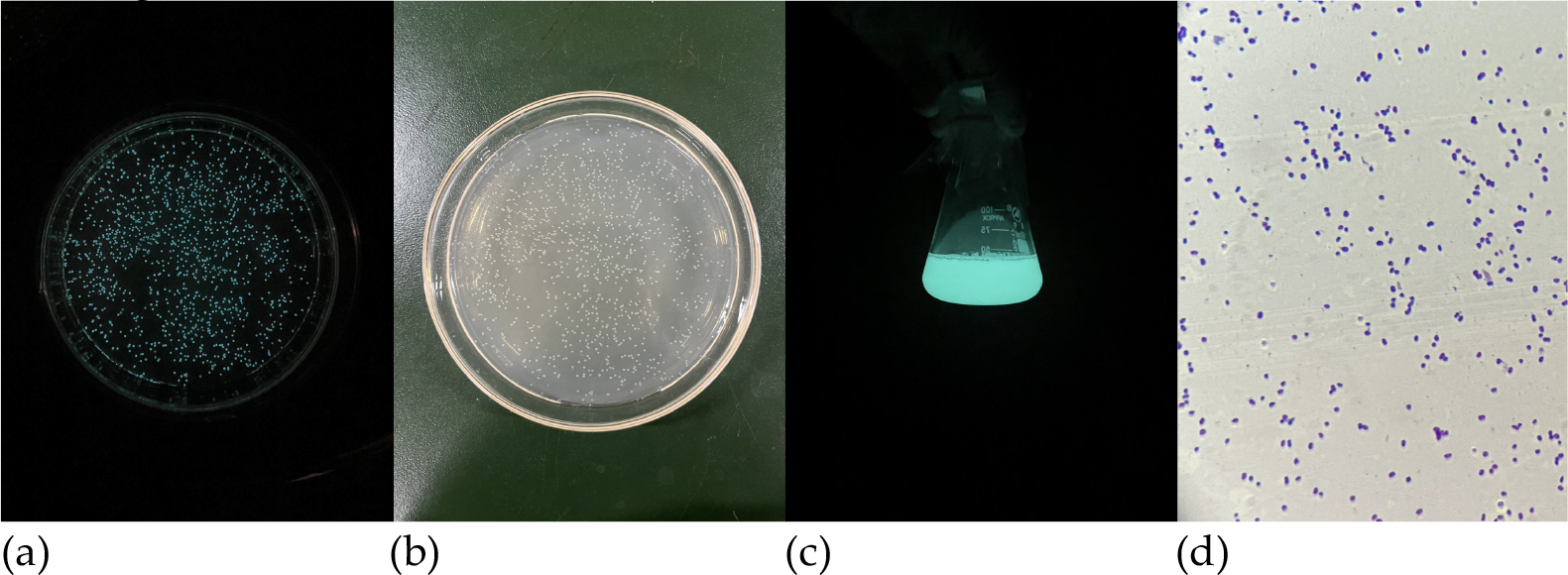
Growth morphology of the strain. (a) Colony morphology on solid medium; (b) Colony morphology on solid medium under dark conditions; (c) Colony morphology on liquid culture medium under dark conditions; (d) Morphology of strains under oil microscope.

#### 3.1.2. 16S rRNA analysis

Afte the 16S rRNA sequence was compared with NCBI database by Blast. The results showed that strain FJ21 belonged to Proteobacteria, Gammaproteobacteria, Vibrionales, Vibrionaceae and *Photobacterium*. Based on the phylogeny of 16S rRNA gene sequence, FJ21clustered with the strain *Photobacterium phosphoreum and Photobacterium kishitanii*.(Fig 2).

**Figure 2.**
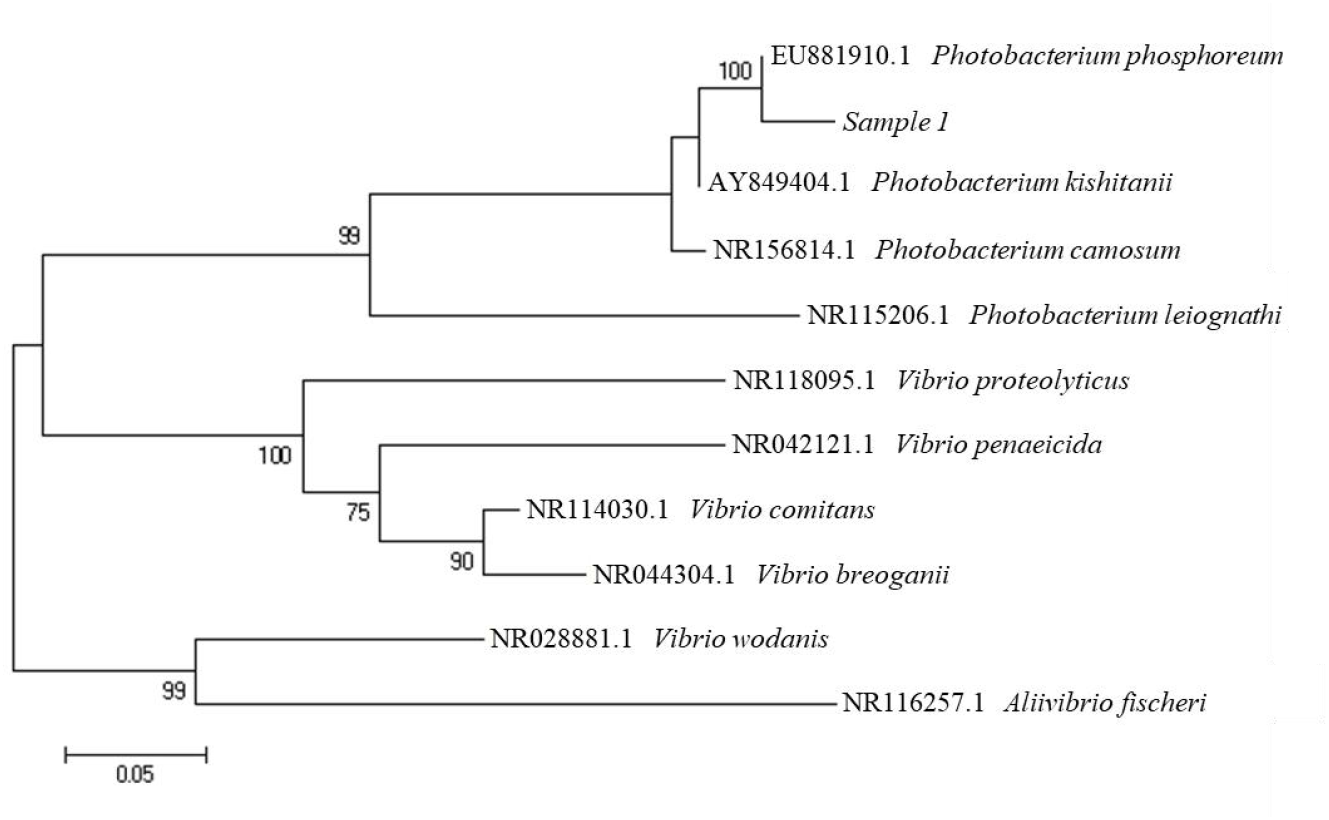
Neighbour Joining phylogenetic tree based on 16S rRNA gene sequences.

#### 3.1.3. Physicochemical properties of the protein encoded by *lux* gene

The sequencing results showed that *lux*C(1437bp), *lux*D(921bp), *lux*A(1074bp), *lux*B(987bp), *lux*F (696bp), *lux*E (1122bp), *lux*G(705bp) genes existed in strain FJ21. Online software Expasy (https://www.expasy.org/) was used to predict the physicochemical properties of proteins encoded by *lux*C, *lux*D, *lux*A, *lux*B, *lux*F, *lux*E, and *lux*G genes (Table 1).

**Table 1.**
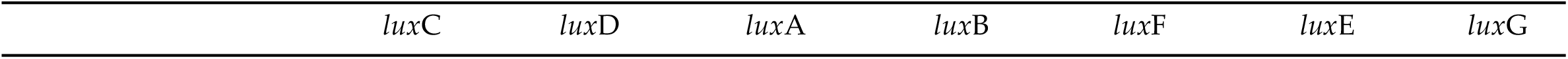
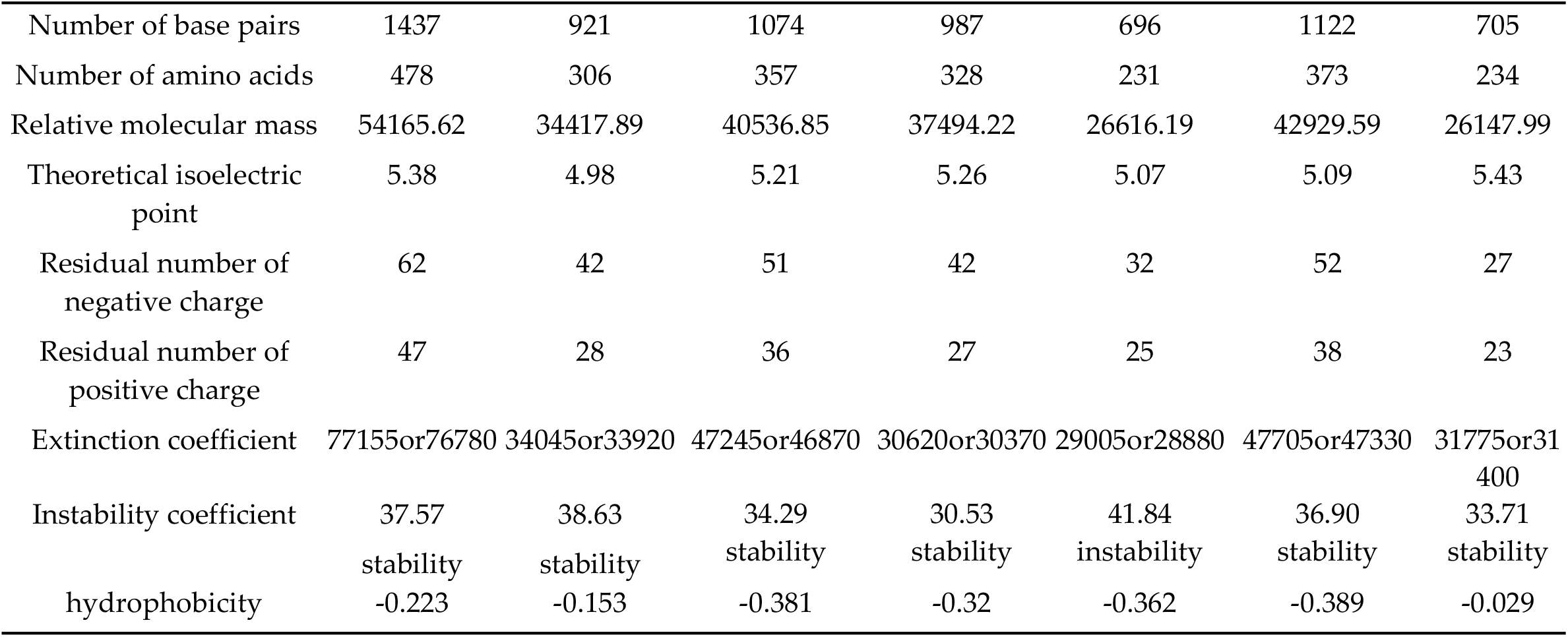
Prediction of physical and chemical properties of the protein encoded by *lux* genes.

It can be seen that the positive charge residues carried by the protein encoded by the gene are less than the negative charge residues, and the isoelectric point is between 4.98 and 5.43, indicating that the protein is easy to precipitate between these values. Only the protein encoded by the *lux*F gene is unstable, and the protein encoded by other genes is stable. In addition, the hydrophobicity is negative, indicating that these proteins have certain hydrophilicity.

#### 3.1.4. Prediction of secondary and tertiary structures proteins encoded by *lux* genes

For unknown proteins, their secondary and tertiary structures can be predicted by amino acid sequences. The online analysis software PSIPRED was used to predict the secondary structure of the protein encoded by *lux* genes (Figure 3). The amino acid sequence of the pink part was α-helix, and the amino acid sequence of the yellow part was β -sheet. SWISS-MODEL database (http://swissmodel.expasy.org/repository/) was used to predict the tertiary structure of proteins (Figure 4). The protein structure in this database was predicted by the homology modeling method. When the sequence similarity between the predicted protein and the template protein exceeds 30 %, the homology modeling method can generate the tertiary structure of the protein with a prediction accuracy of 90%. Except for the *lux* G gene, the similarity of sequences encoded by other genes was more than 30 % after alignment, so the tertiary structure of the protein was closer to the real structure. From the predicted secondary and tertiary structures of *lux* genes encoding proteins, it can be seen that the α subunit and β subunit of the luciferase encoded by *lux*A and *lux*B genes have β-sheet barrel structures, which may be these two genes play an important role in the luminescence activity of luminescent bacteria. Understanding the tertiary structure of proteins is of great significance for studying functional structures (such as molecular docking).

**Figure 3.**
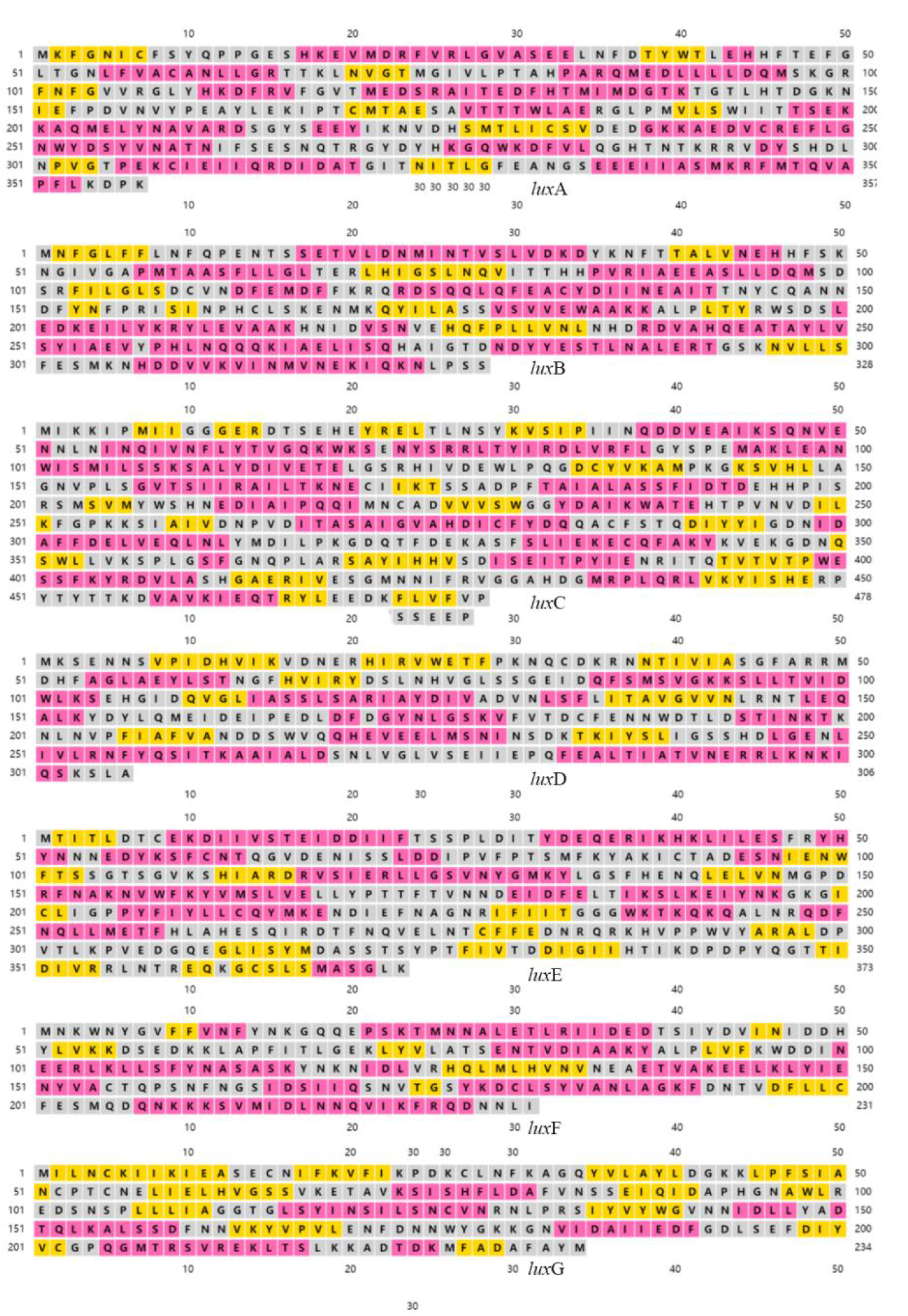
Prediction of protein secondary structure encoded by *lux* genes. The amino acid sequence of the pink part was α-helix, and the amino acid sequence of the yellow part was β-sheet.

**Figure 4.**
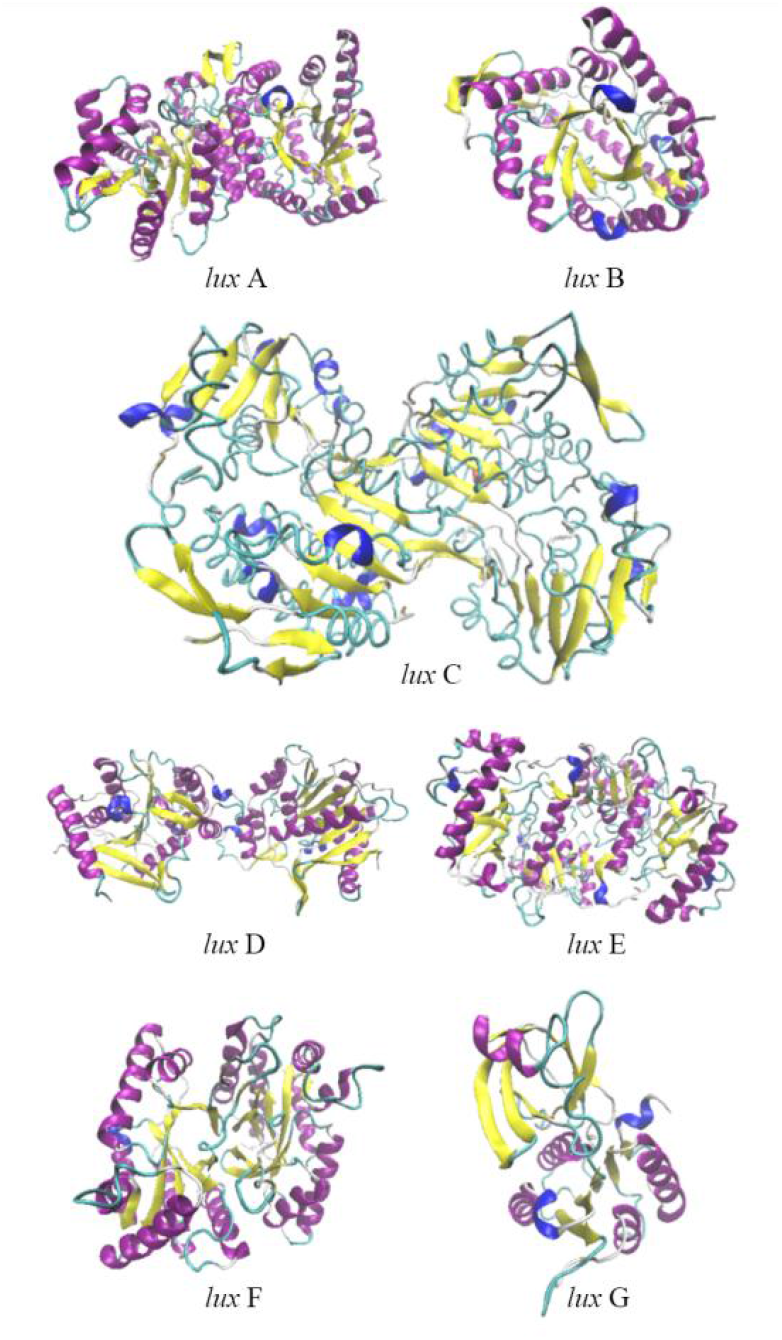
Prediction of protein tertiary structure encoded by *lux* genes, including *lux*A, *lux*B, *lux*C, *lux*D, *lux*E, *lux*F, *lux*G, α-helix (purple), β-sheet(yellow), turn (blue) and random coil(green).

### 3.2. Genome Sequencing and Assembly

The sequencing data are shown in Supplementary Table S1 and the assembly results are shown in Table 2. The genome size was 4853277 bp, the number of coding genes was 4131, and the N50 was 3252201 bp. ATGC content accounted for 30.49 %, 30.29 %, 19.72 %, and 19.50 % of the total base, respectively, and the GC content was 39.23 % (Supplementary Table S2). The genome circle is shown in Figure 5. The genome sequence of strain FJ21 has been submitted to the GenBank database with accession number SRX10356131.

**Table 2.** Assembly result statistics.

| Feature | Data |
| --- | --- |
| Total bases | 4853277 |
| Contig number | 2 |
| Contig N50 | 3252201 |
| Longest Contig | 3252201 |
| Shortest Contig | 1601076 |

**Figure 5.**
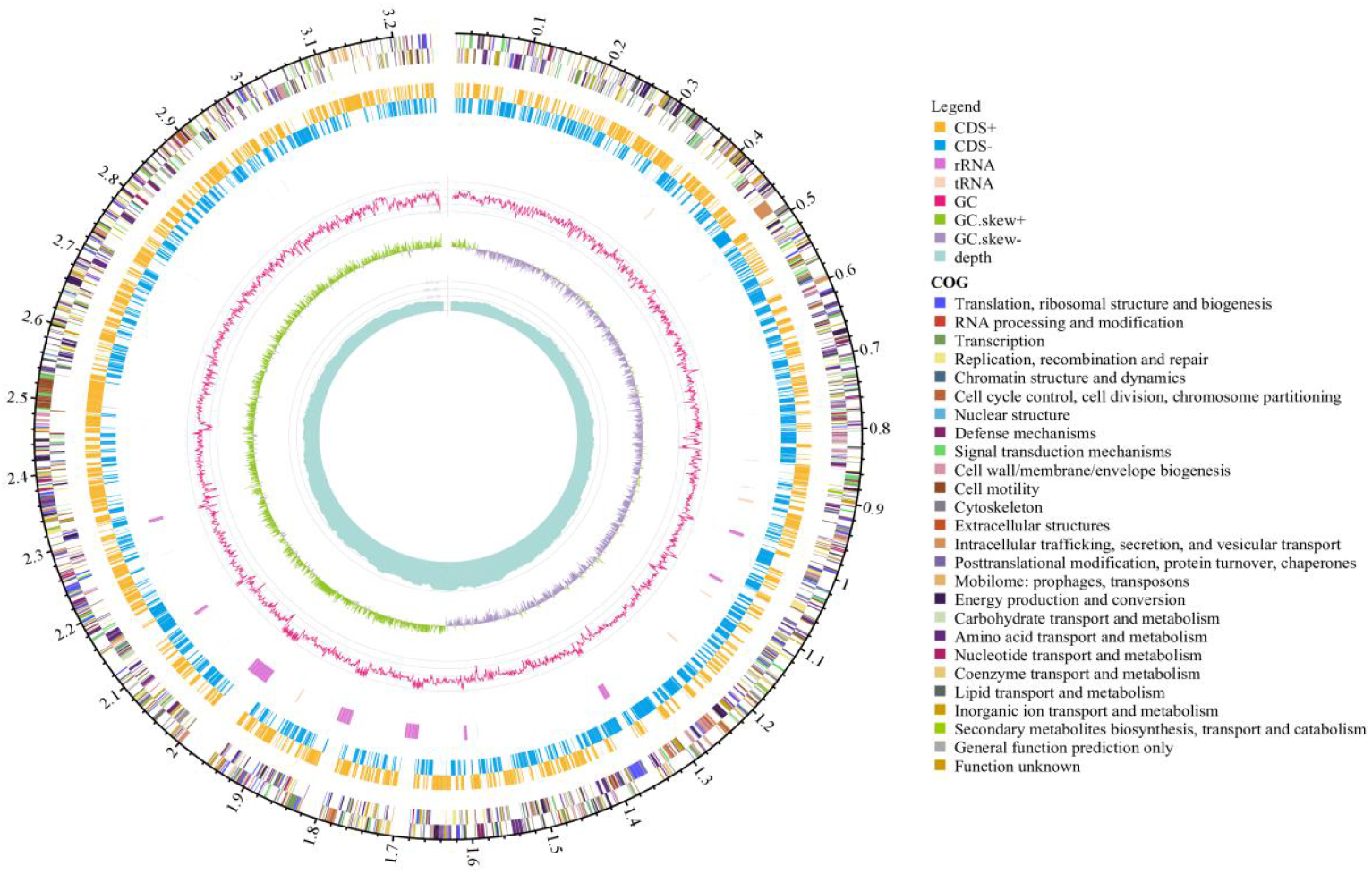
Genome circle diagram of strain FJ21. The characteristics of the marker are shown from outside to inside as follows: colored on clusters of orthologous groups (COG) functional categories, on the forward strand, coding sequences (CDSs); tRNA and rRNA; GC content (outward plots as positive values and inward plots as negative values); GC skew (G - /G + C, the leading chain and the lagging chain can be judged by the change of GC skew, generally the leading chain GC skew > 0, the lagging chain GC skew < 0) and Sequencing depth.

### 3.3. Genome Structure and function Annotation

The genome contains 4131 CDSs, 4027962 bp in length, and a CRISPR sequence (a cluster of regularly spaced short palindromic repeats, often found in many bacteria and archaea). Gene islands are not predicted on the genome. The results of genome structure prediction are shown in Supplementary Table S3.

There are 1,769, 3,141, 2472, 4070, 3514, and 1413 genes that were annotated respectively relatethe d to KEGG pathway, COG category, GO, Refseq, Pfam and TIGRFAMs databases, and annotated results are shown in Supplementary Table S4.

#### 3.3.1. COG function classification

In the COG category(Figure 6), there are 232 Energy production and conversion, 328 Amino acid transport, and metabolism, 210 Carbohydrate Transport and metabolism, 265 Translation, ribosomal structure and biogenesis, 266 Transcription, 267 Cell wall/membrane/envelope biogenesis and 194 Inorganic ion transport and metabolism. The corresponding protein sequence was compared with the COG database to complete the annotation classification of homologous genes, and the coding genes including information storage and processing, cell biology process and signal transduction, basic metabolism, and unknown functions were obtained[35]. As shown in Figure 3, a total of 3141 proteins obtained COG functional annotations, accounting for 76.03 % of the total number of predicted genes, including Energy production and conversion, Amino acid transport and metabolism, Carbohydrate Transport and metabolism, Translation/ribosomal structure and biogenesis, Transcription, Cell wall/membrane/envelope biogenesis, Inorganic ion transport and metabolism, and the number of genes was 232, 328, 210, 265, 266, 267 and 194, respectively.

**Figure 6.**
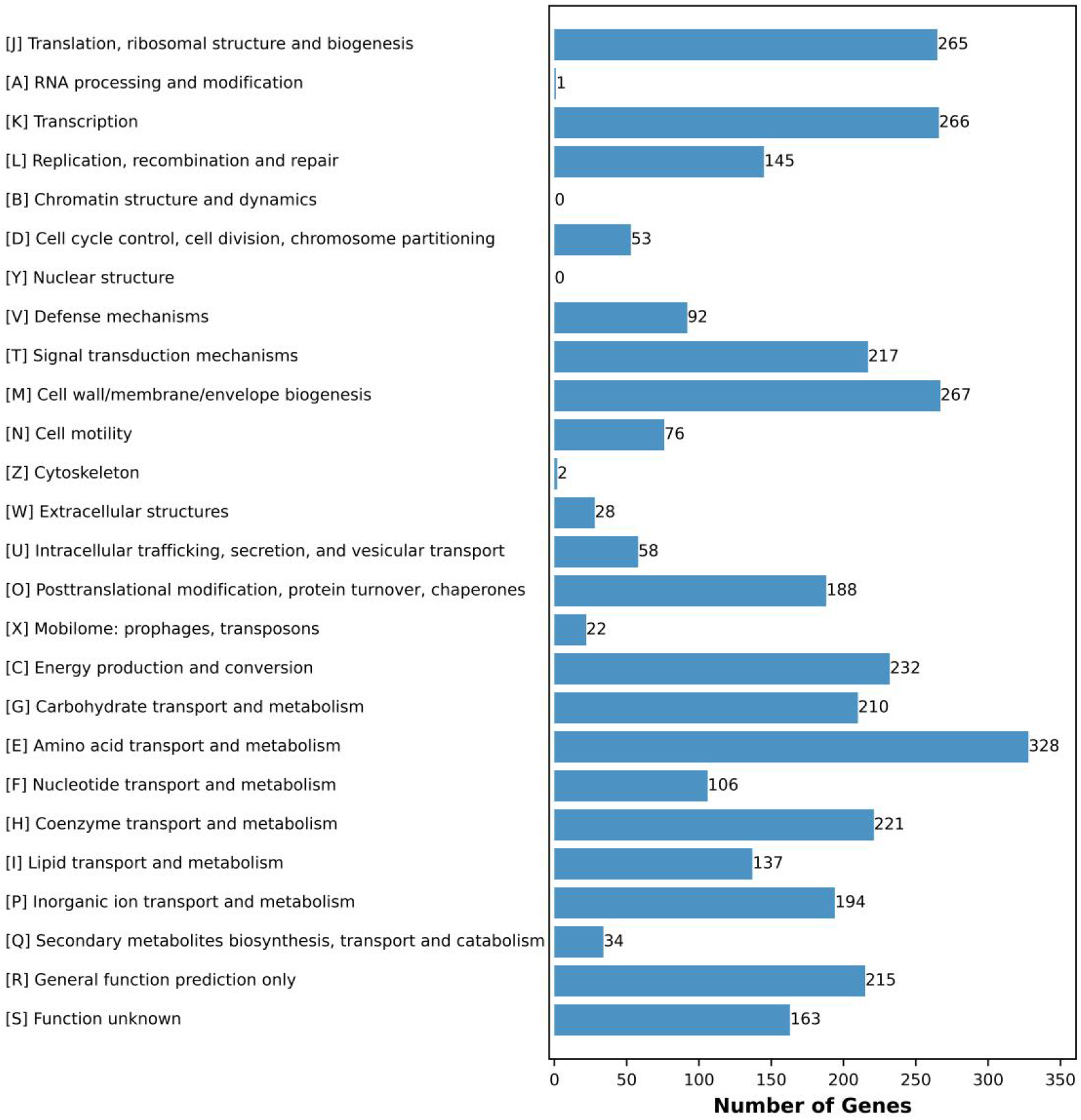
COG functional classification of strain FJ21, including Translation, ribosomal structure and biogenesis, RNA processing and modification, Transcription, Replication, recombination and repair, Cell cycle control, cell division, chromosome partitioning, Defense mechanisms, Signal transduction mechanisms, Cell wall/membrane/envelope biogenesis, Cell motility, Cytoskeleton, Extracellular structures, Intracellular trafficking, secretion, and vesicular transport, Posttranslational modification, protein turnover, chaperones, Mobilome: prophages, transposons, Energy production and conversion, Carbohydrate Transport and metabolism, Amino acid transport and metabolism, Nucleotide transport and metabolism, Coenzyme transport and metabolism, Lipid transport and metabolism, Inorganic ion transport and metabolism, Secondary metabolites biosynthesis, transport and catabolism.

#### 3.3.2. GO function classification

As shown in Figure 7, a total of 2472 genes were annotated for GO function, accounting for 59.84 % of the total number of predicted genes. GO function mainly divides them into molecular function, biological process and cellular component[36]. In molecular function, there are many genes annotated by molecular transducer activity, antioxidant activity and transporter activity. In the biological process, there are many genes annotated by metabolic process, positive regulation of the biological process, and negative regulation of the biological process. In cell components, there are many genes annotated by cell, cell part and membrane part. Therefore, GO functional annotation is more convenient for us to understand the biological significance behind genes.

**Figure 7.**
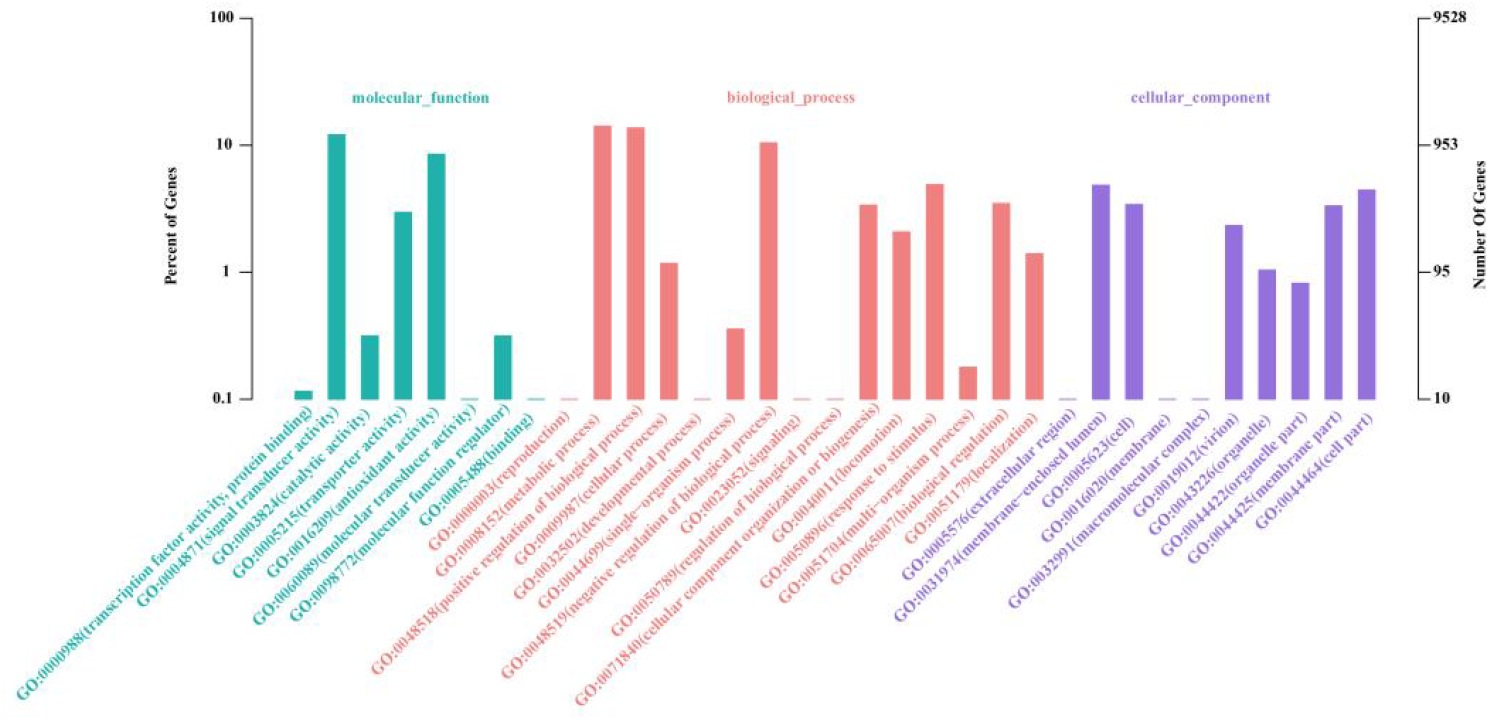
Go function classification diagram. There are three main sections: molecular function(including transcription factor activity, protein binding, signal transducer activity, catalytic activity, transporter activity, antioxidant activity, molecular transducer activity, molecular function regulator, binding), biological process(including reproduction, metabolic process, positive regulation of biological process, cellular process, developmental process, single-organism process, negative regulation of the biological process, signaling, regulation of biological process, cellular component organization or biogenesis, locomotion, response to stimulus, multi-organism process, biological regulation, localization) and cellular component(including extracellular region, membrane-enclosed lumen, cell, membrane, macromolecular complex, virion, organelle, organelle part, membrane part, cell part)

#### 3.3.3. KEGG pathway analysis

The 2945 genes in the KEGG pathway were enriched in 208 metabolic pathways (figure 8), and the number of effectively annotated genes was 1769, accounting for 42.82 % of the total predicted genes. There are five categories, namely Metabolism, Genetic Information Processing, Environmental Information Processing, Cellular processes and Organismal Systems. The most annotated genes in metabolism are carbohydrate metabolism, energy metabolism, and amino acid metabolism. The main pathways are oxidative phosphorylation pathway (ko00190) (40 genes), arginine and proline pathway (ko00330) (16 genes), glycolysis/ gluconeogenesis pathway (ko00010) (31 genes), citric acid cycle (TCA cycle) (ko00020) (19 genes). The least genes were annotated in organismal systems.

**Figure 8.**
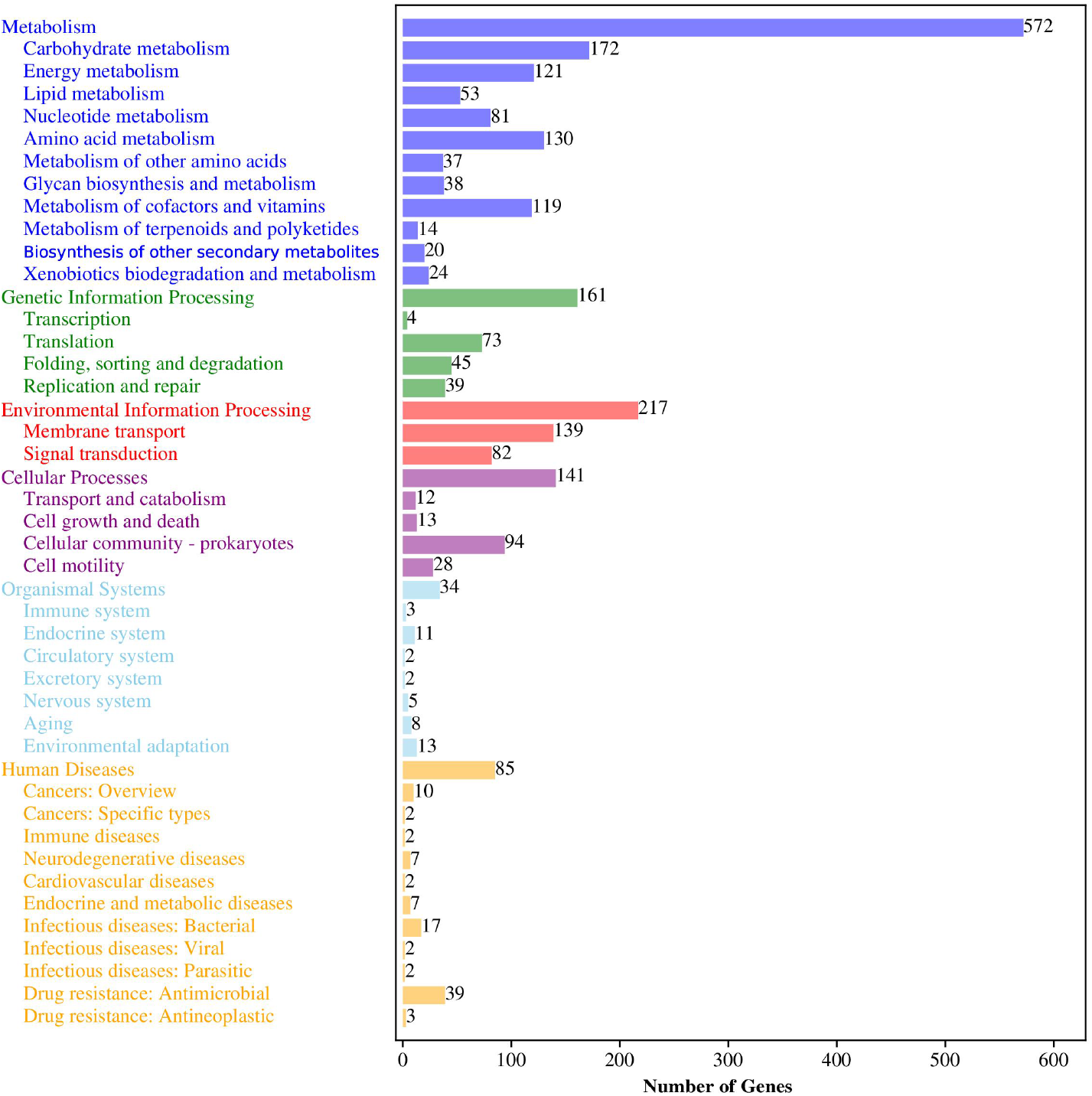
KEGG function classification diagram, including metabolism, genetic information processing, environmental information processing, cellular processes, organismal systems, human diseases; the number of genes (right).

#### 3.3.4. Analysis of interaction genes between pathogen and host

Most luminous bacteria are nonpathogenic[38], while two subspecies of *Vibrio harveyi* and *Photobacterium damselae* are pathogens of many aquatic organisms[37,38]. 870 genes were annotated in the PHI database (Figure 9), of which 326 genes (55.72 %) resulted in reduced virulence. There were 43 increased virulence genes, 217 unaffected pathogenicity genes, 82 loss of pathogenicity genes, 103 effector genand es, resistance to chemical and sensitivity to chemical genes were the least. In this annotation, most of the genes belonged to the reduced virulence genes and unaffected pathogenicity genes. Effector genes are associated with pathogenicity, but increased virulence genes are the key genes.

**Figure 9.**
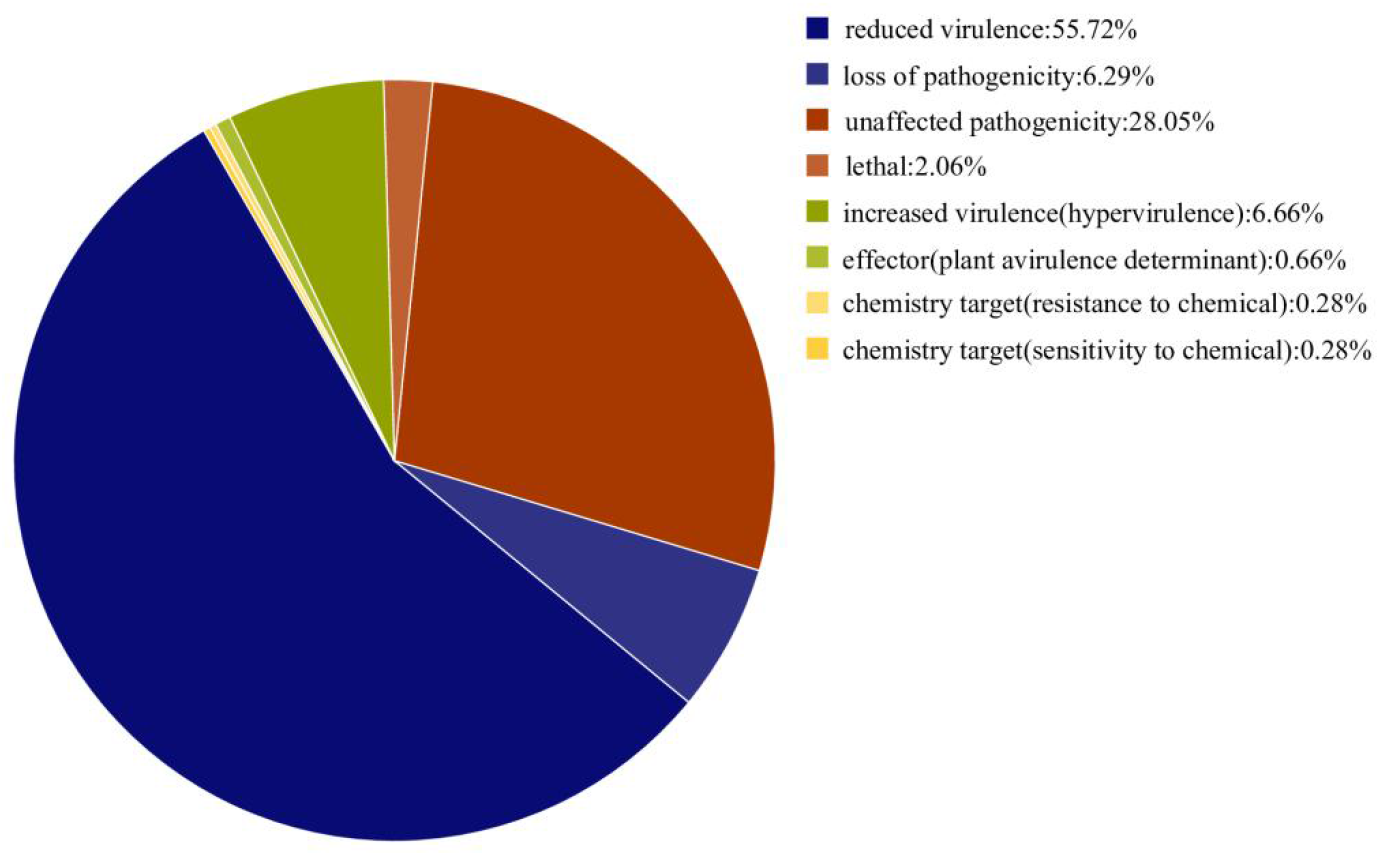
Gene analysis of pathogen host interaction. From left (blue) to right (yellow) as follows: reduced virulence genes; loss of pathogenicity genes; unaffected pathogenicity genes; lethal genes; increased virulence genes; effector genes; chemistry target (resistance to chemical) and chemistry target (sensitivity to chemical) genes.

#### 3.3.5. Annotation of Resistance Genes in the CARD Database

The antibiotic resistance genes were annotated using the CARD database, and the information was shown in Table 3. Including the classification of ARO, the Identities, the classification of antibiotics, the resistance mechanism, and the classification of the AMR gene family. The highest identities can reach 100 %. The resistance mechanisms are antibiotic efflux and antibiotic target Alteration (Supplementary Table S5).

**Table 3.** Antibiotic resistance gene annotation of strain FJ21.

| ARO | %Identities | Drug Class | Resistance Mechanism | AMR Gene Family |
| --- | --- | --- | --- | --- |
| rsmA | 89.29 | A fluoroquinolone antibiotic,<br>diaminopyrimidine antibiotic,<br>phenicol antibiotic | antibiotic efflux | resistance-nodulation-<br>cell division (RND) |
| vanWB | 100 | glycopeptide antibiotic | antibiotic target<br>Alteration | antibiotic efflux<br>pumpvanW,<br>glycopeptide |
| CRP | 94.29 | macrolide antibiotic,<br>fluoroquinolone antibiotic;<br>penam | antibiotic efflux | resistance gene cluster<br>resistance-nodulation-<br>cell division (RND)<br>antibiotic efflux pump |
| Haemophilus<br>influenzae PBP3<br>conferring<br>resistance to beta-lactam antibiotics | 46.85 | cephalosporin; cephamycin;<br>penam | antibiotic target<br>alteration | Penicillin-binding<br>protein mutations<br>conferring resistance to<br>beta-lactam antibiotics |

#### 3.3.6. Analysis of carbohydrate-related enzymes (CAZy)

Carbohydrates are the main source of energy needed to maintain life activities and are the most widely distributed organic compounds in nature. The carbohydrate-related enzymes (CAZy) database collected six categories of enzymes, namely glycoside hydrolases (GHs), glycosyltransferases (GTs), carbohydrate esterases (CEs), carbohydrate-binding modules (CBMs), auxiliary module enzymes (AAs) and polysaccharide lyases (PLs)[39].

In this database, strain FJ21 contains 51 carbohydrate-related enzymes. Among them, glycoside hydrolases (GHs) gene annotation is the most. There are 19 types of 43 genes, accounting for 34.4%. Glycosides produced by enzymatic hydrolysis of glycosidic bonds can be used in biological metabolic pathways. Glycosyltransferases (GTs) have 12 types and 32 genes, accounting for 25.6 %. GTs can participate in a variety of life activities in cells, transferring monosaccharides of active substances in vivo to proteins, lipids, sugars, and nucleic acids to form glycosylation. Carbohydrate esterases (CEs) and carbohydrate-binding modules (CBMs) accounted for 16 %, respectively. The auxiliary modular enzymes (AAs) and polysaccharide lyases (PLs) genes were the least.

### 3.4. Phylogenetic tree, ANI and Tetra analysis

#### 3.4.1. Genome phylogenetic tree analysis

Then the whole genome sequence(Supplementary Table S6) was constructed a phylogenetic tree (Figure 10). The results showed that *Photobacterium kishitanii* ATCCBAA-1194, *Photobacterium phosphoreum* JCM21184, *Photobacterium aquimaris* LC2-065, *Photobacterium malacitanum* CECT9190, and *Photobacterium carnosum* TMW 2.2021 were clustered together in the phylogenetic tree. We found that the whole genome sequence of strain FJ21 showed a great deal of similarity with the genome of *Photobacterium kishitanii* ATCCBAA-1194.

**Figure 10.**
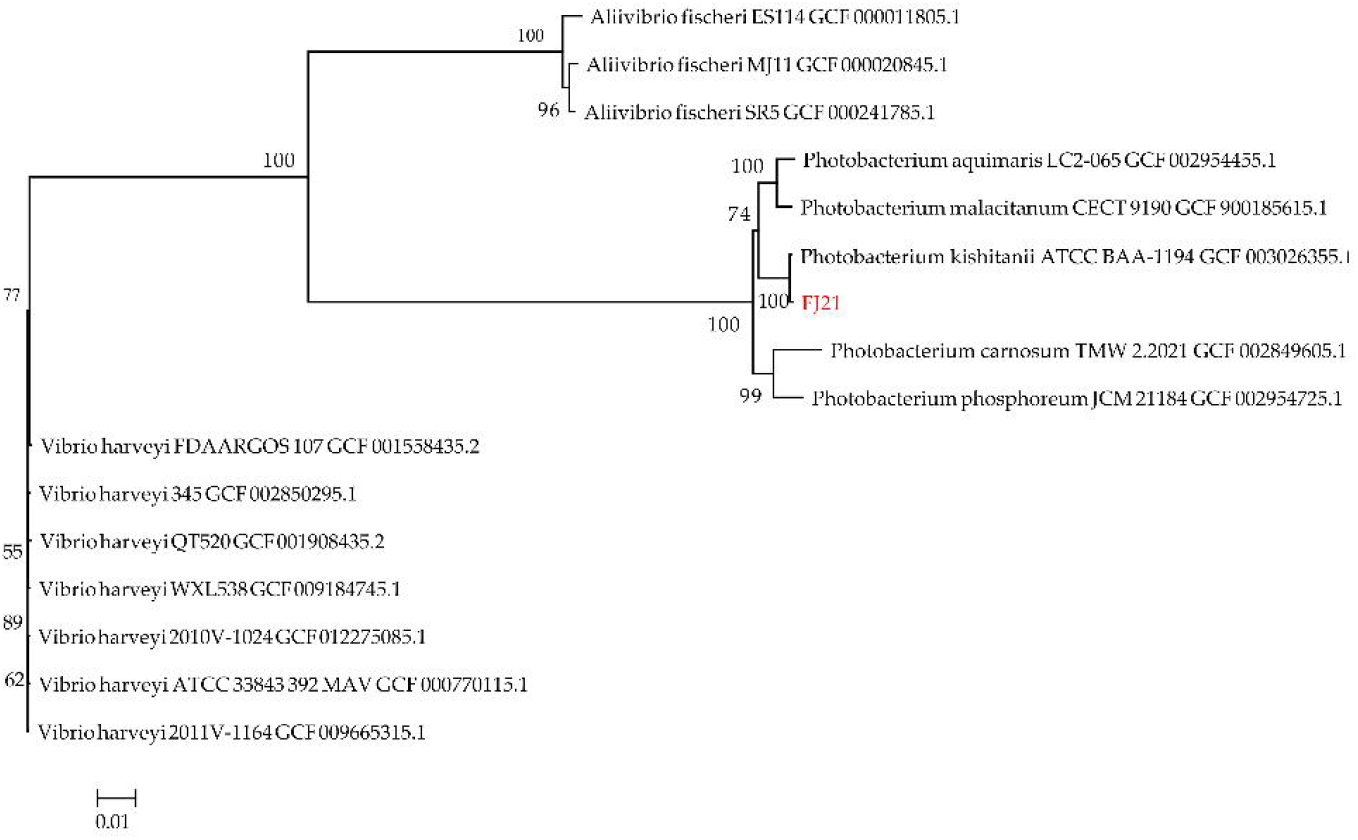
Phylogenetic tree based on whole-genome genes sequences.

The difference between the two results may be due to the effect of gene transfer.

#### 3.4.2. ANI and TETRA analysis

Average nucleotide identity (ANI) and tetra nucleotide signatures (TETRA) between strain FJ21 and different *Photobacterium* strains were calculated (Table 4). The ANI value of strain FJ21 against *Photobacterium kishitanii* were 97.61% (ANIb, based on BLAST) and 97.55% (ANIm, based on MUMmer), respectively. Both were higher than the defined threshold (95%). In contrast, against the others, the ANI value was down to 84.68-87.32%, indicating strain FJ21 was phylogenetically close to *Photobacterium kishitanii*. Therefore, strain FJ21 should be reclassified as *Photobacterium kishitanii* rather than *Vibrio fischeri* or *Photobacterium phosphoreum*. The results of Tetra also supported the conclusion.

**Table 4.** ANI and tetra nucleotide signatures (TETRA) analyses between strain FJ21 and *Photobacterium* strains.

| Species name | FJ21 |  |  | *P.kishitanii* |  |  | *P.phosphoreum* |  |  | *P.aquimaris* |  |  | *P.malacitanum* |  |  | *P.carnosum* |  |  |
| --- | --- | --- | --- | --- | --- | --- | --- | --- | --- | --- | --- | --- | --- | --- | --- | --- | --- | --- |
|  | AN lb | AN Im | Tet ra | AN lb | AN Im | Tet ra | AN lb | AN Im | Tet ra | AN lb | AN Im | Tet ra | AN lb | AN Im | Tet ra | AN lb | AN Im | Tet ra |
| FJ21 | * | * | * | 97.55 | 97.81 | 0.998 | 85.68 | 87.32 | 0.985 | 85.03 | 87.17 | 0.993 | 84.72 | 86.90 | 0.992 | 84.62 | 86.68 | 0.988 |
| *P.kishitanii* | 97.61 | 97.80 | 0.998 | * | * | * | 85.41 | 87.32 | 0.988 | 84.82 | 87.13 | 0.994 | 84.41 | 86.86 | 0.993 | 84.23 | 86.72 | 0.990 |
| *P.phosphoreum* | 85.64 | 87.32 | 0.985 | 85.64 | 87.32 | 0.988 | * | * | * | 84.89 | 86.86 | 0.991 | 84.48 | 86.57 | 0.990 | 85.66 | 87.69 | 0.992 |
| *P.aquimaris* | 85.11 | 87.17 | 0.993 | 85.16 | 87.13 | 0.994 | 85.03 | 86.86 | 0.991 | * | * | * | 92.61 | 92.97 | 0.998 | 84.10 | 86.87 | 0.989 |
| *P.malacitanum* | 84.68 | 86.9 | 0.992 | 84.62 | 86.86 | 0.993 | 84.56 | 86.57 | 0.990 | 92.44 | 92.96 | 0.998 | * | * | * | 83.23 | 86.09 | 0.986 |
| *P.carnosum* | 84.68 | 86.69 | 0.988 | 84.65 | 86.72 | 0.990 | 86.07 | 87.69 | 0.992 | 84.26 | 86.88 | 0.989 | 83.42 | 86.10 | 0.986 | * | * | * |

### 3.5. Genome collinearity analysis

#### 3.5.1. The basic characteristics of genomes

The genome size of the six strains is similar, ranging from 4380538bp to 4853277bp, and the number of coding genes is 3739-4131 (Table 5). The genome characteristics of different strains of the same bacteria are closer, the stain FJ21 is closer to *Photobacterium kishitanii* in genome size. Compared with the number of coding genes and RNA of other strains, the results showed that the number of coding genes and RNA predicted were significantly higher than others.

**Table 5.** Genome features of *P.kishitanii, P.phosphoreum, P.aquimaris, P.malacitanum, P.carnosum* and strain FJ21.

| Species name | Genome size (bp) | Coding gene | tRNAs | 5S、16S、23S rRNA |
| --- | --- | --- | --- | --- |
| *P.kishitanii* | 4732354 | 4087 | 159 | 6, 6, 5 |
| *P.phosphoreum* | 4550107 | 3840 | 73 | 3, 0, 1 |
| *P.aquimaris* | 4525475 | 3826 | 83 | 1, 0, 1 |
| *P.malacitanum* | 4380538 | 3739 | 186 | 10, 15, 12 |
| *P.carnosum* | 4559543 | 3984 | 122 | 8, 1, 8 |
| FJ21 | 4853277 | 4131 | 226 | 24, 22, 22 |

#### 3.5.2. Collinearity analysis

MUMmer (version 3.23) software was used to compare the strain FJ21 with *Photobacterium kishitanii* ATCCBAA-1194, *Photobacterium phosphoreum* JCM21184, *Photobacterium aquimaris* LC2-065, *Photobacterium malacitanum* CECT9190, and *Photobacterium carnosum* TMW 2.2021. The collinearity and structural variation of genomic sequences are shown in Figure 11, and there were 218, 744, 717, 748, and 708 contrast blocks between *Photobacterium kishitanii* ATCCBAA-1194, *Photobacterium phosphoreum* JCM21184, *Photobacterium aquimaris* LC2-065, *Photobacterium malacitanum* CECT9190, and *Photobacterium carnosum* TMW 2.2021 and the strain FJ21, respectively. They accounted for 86.92 %, 70.27 %, 64.82 %, 65.16 % and 56.32 % of the genome of the strain, respectively. According to the results, the collinearity between genomes is good, but there are a small number of genome rearrangement events such as inversion and translocation. It can be seen that the six strains still have great differences in evolution.

**Figure 11.**
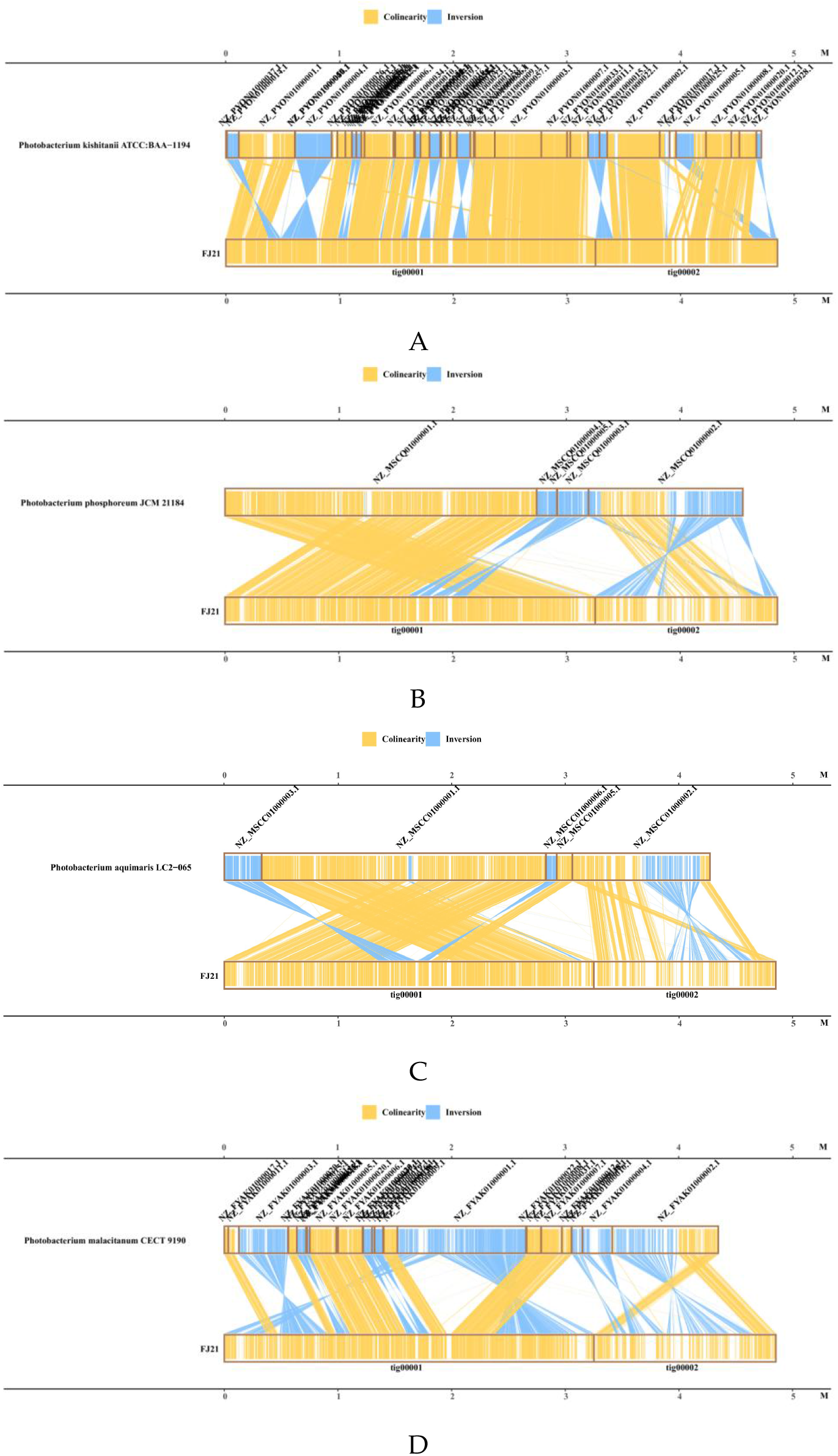
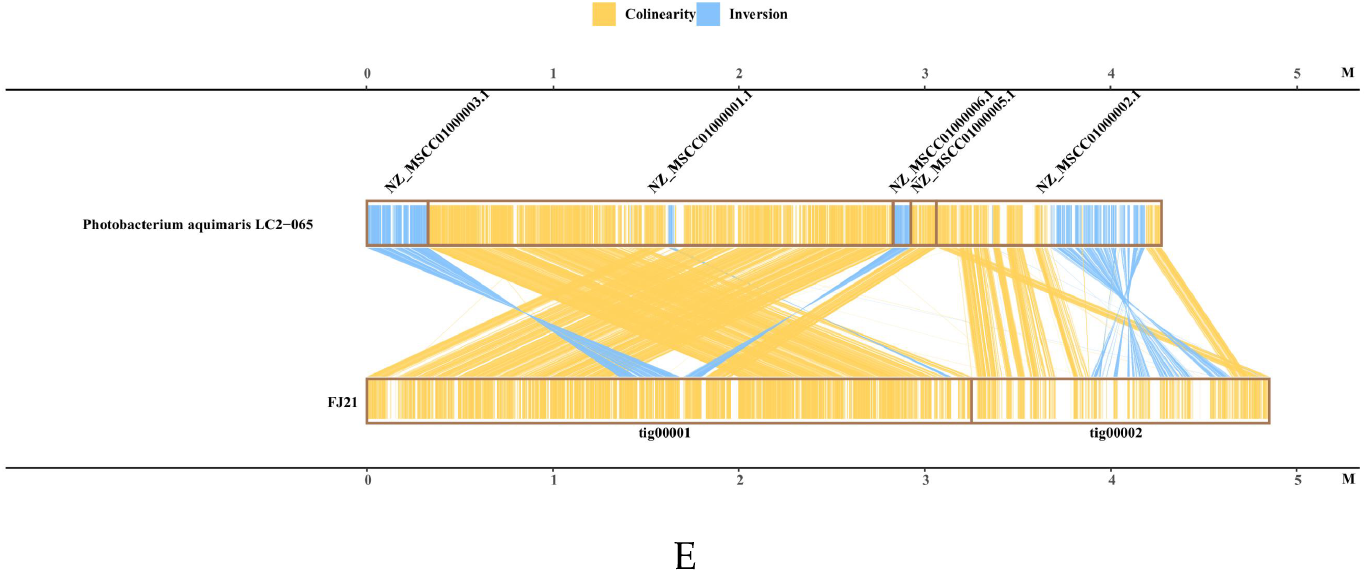
Collinearity analysis of strain FJ21, *P. kishitanii, P. phosphoreum, P. aquimaris, P. malacitanum* and *P. carnosum*. (A) Collinearity analysis of *P. kishitanii* and strain FJ21, (B) Collinearity analysis of *P. phosphoreum* and strain FJ21, (C) Collinearity analysis of *P. aquimaris* and strain FJ21, (D) Collinearity analysis of *P. malacitanum* and strain FJ21, (E) Collinearity analysis of *P. carnosum* and strain FJ21.

### 3.6. Secondary metabolite gene clusters

The encoding genes of secondary metabolites are usually clustered in the genome, encoding complex enzymes with multiple functions. AntiSMASH (version 6.0.0) software was used to predict the gene cluster of the assembled genome. Three types of secondary metabolite gene clusters(RiPP-like, beta lactone, and arylpolyene) were predicted in the FJ21 genome(Table 6), and siderophore only exists in *Photobacterium phosphoreum*.

**Table 6.**
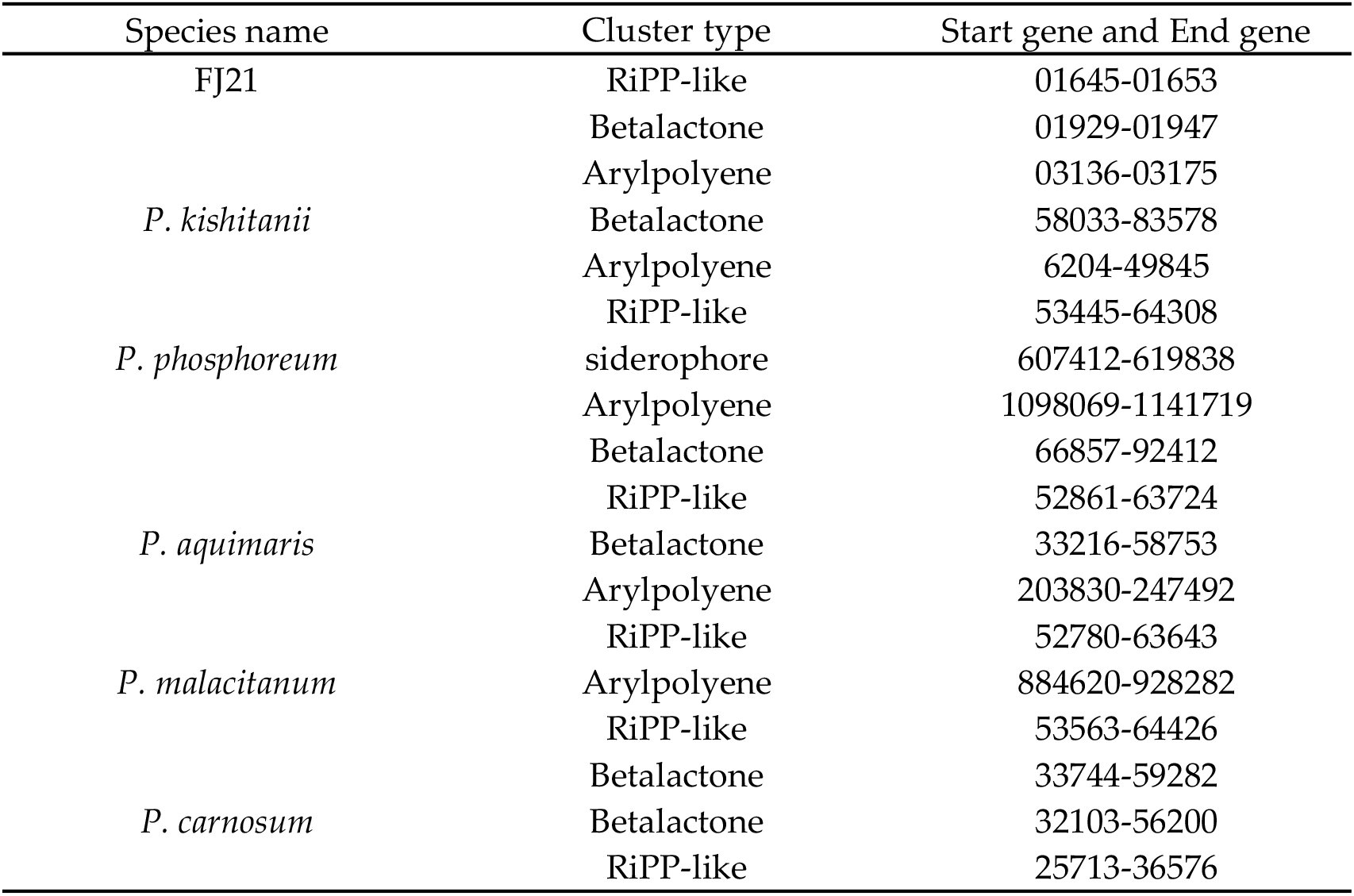
Gene clusters of a secondary metabolite of *Photobacterium kishitanii* ATCCBAA-1194, *Photobacterium phosphoreum* JCM21184, *Photobacterium aquimaris* LC2-065, *Photobacterium malacitanum* CECT9190, and *Photobacterium carnosum* TMW 2.2021 and strain FJ21.

### 3.7. For detection of heavy metal toxicity

The fitted concentration-response curve of the single toxicity of heavy metal to the strain FJ21 is shown in the figure 12, and the parameter values of concentration-dose effect curve are shown in the table 7. With the increase of heavy metal concentration, the inhibition rate of bacteria’s luminescence gradually increased. R^2^ values are all greater than 0.98, which indicates that the concentration of heavy metals is positively related to its inhibition rate of luminescent bacteria. The EC_50_ value can reflect the toxicity of pollutants. The smaller the EC50 value, the greater the toxicity. According to the table, the EC_50_ values of Pb(NO_3_)_2_, ZnSO_4_·7H_2_O, CdCl_2_·2.5H_2_O, CuSO_4_·5H_2_O and K_2_Cr_2_O_7_ to luminescent bacteria are 1.11mg/L, 1.57mg/L, 28.63mg/L, 77.35mg/L and 201.80mg/L, respectively. The toxicity is Pb(NO_3_)_2_ > ZnSO_4_·7H_2_O > CdCl_2_·2.5H_2_O > CuSO_4_·5H_2_O > K_2_Cr_2_O_7_.

**Table 7.** parameter values of concentration-dose effect curve.

| Heavy metal | EC<sub>50</sub>(mg/L) | Ion mass concentration | R<sup>2</sup> |
| --- | --- | --- | --- |
| ZnSO<sub>4</sub>·7H<sub>2</sub>O | 1.57 | 0.36 | 0.9994 |
| CuSO<sub>4</sub>·5H<sub>2</sub>O | 77.35 | 19.80 | 0.9988 |
| Pb(NO<sub>3</sub>)<sub>2</sub> | 1.11 | 14.09 | 0.9962 |
| CdCl<sub>2</sub>·2H<sub>2</sub>O | 28.63 | 0.69 | 0.9848 |
| K<sub>2</sub>Cr<sub>2</sub>O<sub>7</sub> | 201.80 | 35.67 | 0.9992 |

**Figure 12.**
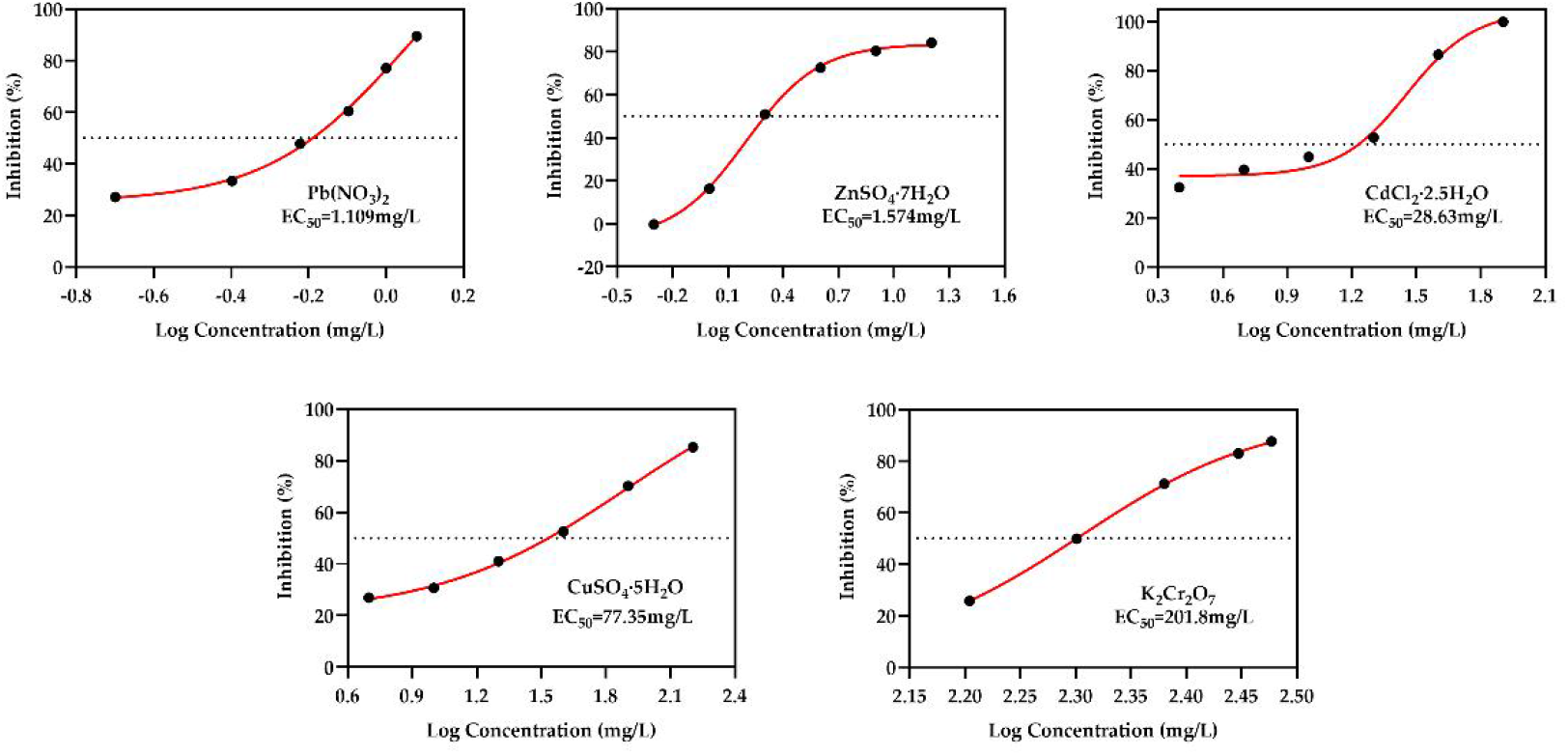
The fitted concentration-response curve of the single toxicity of heavy metal to the strain FJ21. The abscissa represents the logarithm of heavy metal concentration, and the ordinate represents the luminescence inhibition rate of the strain FJ21.

## 4. Conclusions

The colony shape is round, slightly raised, smooth, milky white and short rod, which emits bright blue-green light in the dark. Based on 16S rRNA the strain FJ21 was phylogenetically close to the strain *Photobacterium phosphoreum and Photobacterium kishitanii*. While the whole genome, ANI, TETRA analyses and collinearity analysis further verified the relationship between strain FJ21 and the species of *Photobacterium kishitanii* ATCCBAA-1194. Therefore, we argue that strain FJ21 should be classified as a strain of *Photobacterium kishitanii*. This indicates that the whole genome analytic method is very important for species identification.

The strain contains a chromosome with a total length of 4853277bp and GC content of 39.23%. According to the predicted secondary and tertiary structure of the *lux* gene and its encoded protein, the strain contained *lux*C, *lux*D, *lux*A, *lux*B, *lux*F, *lux*E, and *lux*G genes. However, the function of *lux*F gene is still uncertain. Understanding the *lux* genes will help to understand luminescent activities and the mechanism.

In the toxicity test, the toxicity of heavy metals to strain FJ21 is as follows: Pb(NO_3_)_2_ > ZnSO_4_·7H_2_O > CdCl_2_·2.5H_2_O > CuSO_4_·5H_2_O > K_2_Cr_2_O_7_.

## Supplementary Materials

Table S1: Sequencing data statistics. Table S2: Assembly results statistics. Table S3: Genome structure prediction statistics. Table S4: Functional annotation statistics of genome-encoded proteins. S5: Annotation of Resistance Genes in the CARD database. Table S6: The list of genomes used in the study. Word: the strain FJ21 genomic.

## Data Availability Statement

The available complete genome sequence has been admitted to GenBank databases with accession number SRX10356131.

## Acknowledgments

Authors would like to thank the Hunan Provincial Engineering Research Center of Applied Microbial Resources Development for Livestock and Poultry for technical supporting.

